# SMLMFlow: Improving Structural Resolution in Single Molecule Localization Microscopy with Flow Matching

**DOI:** 10.64898/2026.06.11.731424

**Authors:** Sebastian Bauer, Luca Panconi, Inês Cunha, Emma Latron, Daniel Sage, Ruby Peters, Juliette Griffié

## Abstract

While Single Molecule Localization Microscopy (SMLM) aims to generate precise coordinates of molecular targets in cells, the resulting point clouds are inherently blurred by additive noise sources across the experimental, imaging, and processing workflow. This blurring often limits SMLM’s ability to accurately quantify complex assembled structures required to address biological issues, despite reported localization precision down to a couple of nanometers. Here, we present SMLMFlow, a machine learning framework for improving structural resolution in SMLM datasets that combines a graph neural network and a hierarchical transformer with flow matching. We show that SMLMFlow improves structural resolution and downstream quantification across different structures, including filaments and protein nano-clusters, and generalizes to new unseen photophysics models.

## Introduction

Single-molecule localization microscopy (SMLM) enables fluorescence imaging beyond the diffraction limit by temporally separating the emission of individual fluorophores through stochastic blinking [1; 2]. This produces sparse, well-separated point spread functions (PSFs) collected over thousands of camera frames, from which emitters’ positions can be localized with high precision. SMLM datasets thus differ vastly from conventional pixelated microscopy images in that they consist of lists of coordinates, also called point clouds, representing localized emitters. Whilst a fluorophore can, in principle, be localized with a precision (or uncertainty) of a couple of nm, this does not directly translate into high structural resolution [3]. Indeed, SMLM measurements are affected by several additive noise sources along the data pipeline: sample preparation (e.g., immunostaining), microscopy acquisition (e.g., camera, photophysics), and image reconstruction (e.g., fitting). These, combined with stochastic sampling, limit our ability to accurately resolve precise continuous structures that emerge through the collective organization of molecules, such as fibers and clusters. In these cases, structural resolution cannot be accurately reflected by the localization uncertainty alone. This constraint has been well characterized for nano-clusters, leading for example to a systematic overestimation of cluster size and missed cluster identification [4]. Fibrous structures of the cytoskeleton, such as microtubules and actin, whose biological function depends on filament continuity, orientation, and network connectivity across tens to hundreds of nanometers, are a compelling example of structures whose accurate reconstruction is fundamentally limited by these noise sources. Even when individual localizations are precise (e.g.: uncertainty <15 nm), SMLM point clouds of filamentous structures are typically resulting in fragmented or merged fibers. This explains why fiber tracing remains scarce [5]. Improving structural resolution is therefore critical for extracting biologically meaningful information from SMLM data.

Whilst over the last decade several effective frameworks have been developed to address high PSF density, no existing method specifically addresses the challenge of improving structural resolution. Similarly, SMLM applications in machine learning (ML) have primarily focused on either PSF based localization only [6; 7]) or developed analysis pipelines that systematically convert spatial point patterns (SPPs) into pixelated images [8], resulting in information loss. In contrast to the previous work, here we introduce SMLMFlow, a generative framework that directly operates on raw SMLM localization lists (point clouds) and explicitly models the transformation from noisy point clouds to their ground-truth coherent biological structure. We combine cutting-edge ML methodologies: flow matching [9], a generative framework that learns to transport a noisy distribution toward a clean target, paired with an Erwin transformer [10], which consists of a hierarchical point cloud architecture whose internal graph neural network (GNN) [11] layers capture both local molecular neighborhoods and long-range structural context, making it well-suited to the multi-scale geometry of SMLM data. We apply SMLMFlow to both simulated and experimental SMLM datasets of filaments and nano-clusters. In both cases, SMLMFlow improves structural resolution and enables improved characterization of the nano-scale organization beyond what standard SMLM reconstructions, such as ThunderSTORM [12], provide. We further validate that our model generalizes to novel photophysics models.

## Methods

SMLMFlow integrates a conditional flow matching strategy that learns a vector field to transport samples from an easy-to-sample source distribution to the target data distribution via an ordinary differential equation (ODE) [9], [13]. This enables an efficient deterministic generation, from arbitrary source distributions including non-Gaussian, in contrast with diffusion models [14]. State-of-the-art flow matching methods, however, operate on minimal point representations (e.g., spatial coordinates alone) and assume relatively homogeneous point clouds [15], processing each localization without explicit awareness of the surrounding context, such as local geometry or extended structural patterns. This constraint precludes its direct applicability to SMLM datasets, which are characterized by heterogeneous densities, complex multi-scale molecular organization, and additional features that encode information beyond spatial coordinates (e.g., intensity, frame index and PSF width).

To overcome these limitations, SMLMFlow also incorporates an Erwin transformer [10], a hierarchical architecture that organizes localizations into a nested set of spatial neighborhoods, grouping nearby points into local clusters, then clusters into larger regions. This allows the model to simultaneously capture fine-grained local geometry and long-range structural patterns such as fiber orientation or cluster boundaries (see Online Methods). In this case, the Erwin transformer design enables us to retrieve spatial context at multiple scales, yielding a unique embedding for each point that encodes both local geometry and global structural organization. These learned point-wise representations, in addition to photophysics features, serve as inputs to the flow matching decoder, enabling scalable, structurally context-aware transport of localization coordinates.

SMLMFlow’s overall workflow is summarized in Fig. 1A and Online Methods. SMLMFlow takes as input reconstructed SMLM localization lists that include emitters’ spatial coordinates, the camera frame index, the collected intensity, the localization uncertainty, and the background intensity. This information is readily available regardless of the reconstruction algorithm used. During training, paired noisy and ground truth “clean” SPPs are provided. Because SMLM experiments do not provide access to biological ground-truth molecular coordinates, SMLMFlow cannot be trained directly in a supervised manner on experimental data. Simulated data sets were therefore obtained using vSMLM [16; 17]. This realistic virtual SMLM microscope yields previously described cumulative sources of noise and continuous photophysics transitions at the single fluorophore level (Online Methods). Once trained, SMLMFlow can be used for inference on new, unseen data sets, generating a novel SPP with better-resolved structures.

**Figure 1.**
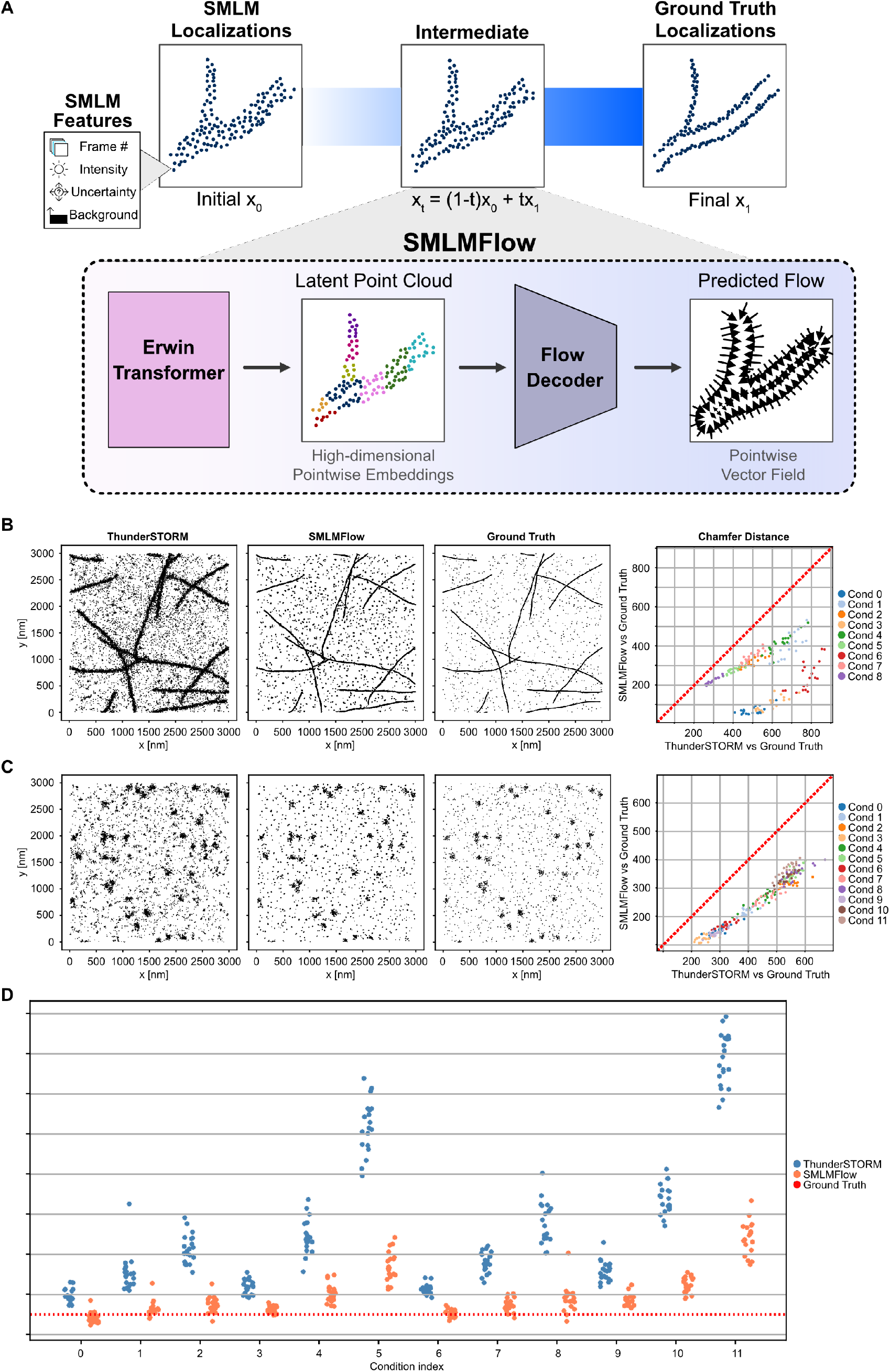
**(A)** SMLMFlow’s overall pipeline. A conditional flow matching model for SMLM point cloud transformation. During training, a source x0 (SMLM localizations from ThunderSTORM) point cloud is interpolated towards the target x1 point cloud to yield *x*_*t*_. Point coordinates and SMLM features (frame index, intensity, uncertainty, background) are processed by an Erwin Transformer, and then a Flow Decoder predicts the flow. **(B)** Left: Example comparison of ThunderSTORM, SMLMFlow, and Ground Truth on a simulated fiber from condition 4. Right: Chamfer Distance evaluation on the simulated STORM fiber dataset. **(C)** as **(B)**, but for the simulated nano-cluster dataset from condition 11. **(D)** Cluster size evaluation of SMLMFlow on the simulated STORM nano-cluster dataset. The average standard deviation of the detected clusters aligns with SMLMFlow closer to the ground truth cluster size (25 nm) than with ThunderSTORM.

## Results

We trained SMLMFlow on simulated fiber- and cluster-like structures spanning a broad range of spatial distributions, randomly positioned within a 3,000 nm× 3,000 nm region of interest (ROI) (Online Methods Fig. 3 & 4). Training data was generated using vSMLM with a three-state STORM photophysics model and *in silico* antibody labeling (see Online Methods and Supp. Figure 5). Localizations were reconstructed using ThunderSTORM [12] in a multi-emitter fitting (MEF) setting, and the dataset was split into training and validation sets (80/20). On validation ROIs, SMLMFlow substantially improved structural resolution compared to raw localizations (Fig. 1B, C, Left). Quantitative evaluation using the one-directional Chamfer distance – the average nearest-neighbor distance from each point in the source to the target point cloud – confirmed significant improvements for both structure types, with average reductions per condition ranging from 24.06% to 86.05% for fiber conditions and 31.74% to 50.37% for cluster conditions (Fig. 1B, C, Right). Downstream analysis further demonstrated improved biological fidelity: cluster descriptors derived from SMLMFlow–processed data, particularly cluster size, more closely matched ground truth across all conditions (Fig. 1D, Online Methods Fig. 6, Online Methods). Evaluation on unseen simulated parallel lines with varying inter-spacing showed that SMLMFlow resolves structures separated by as little as 50 nm, corresponding to a 10 nm gain in effective structural resolution (Fig. 2A). Inference required 1.7 seconds on average for ROIs with an average of 22592 localizations on a GPU NVIDIA A100-SXM4-40GB device.

**Figure 2.**
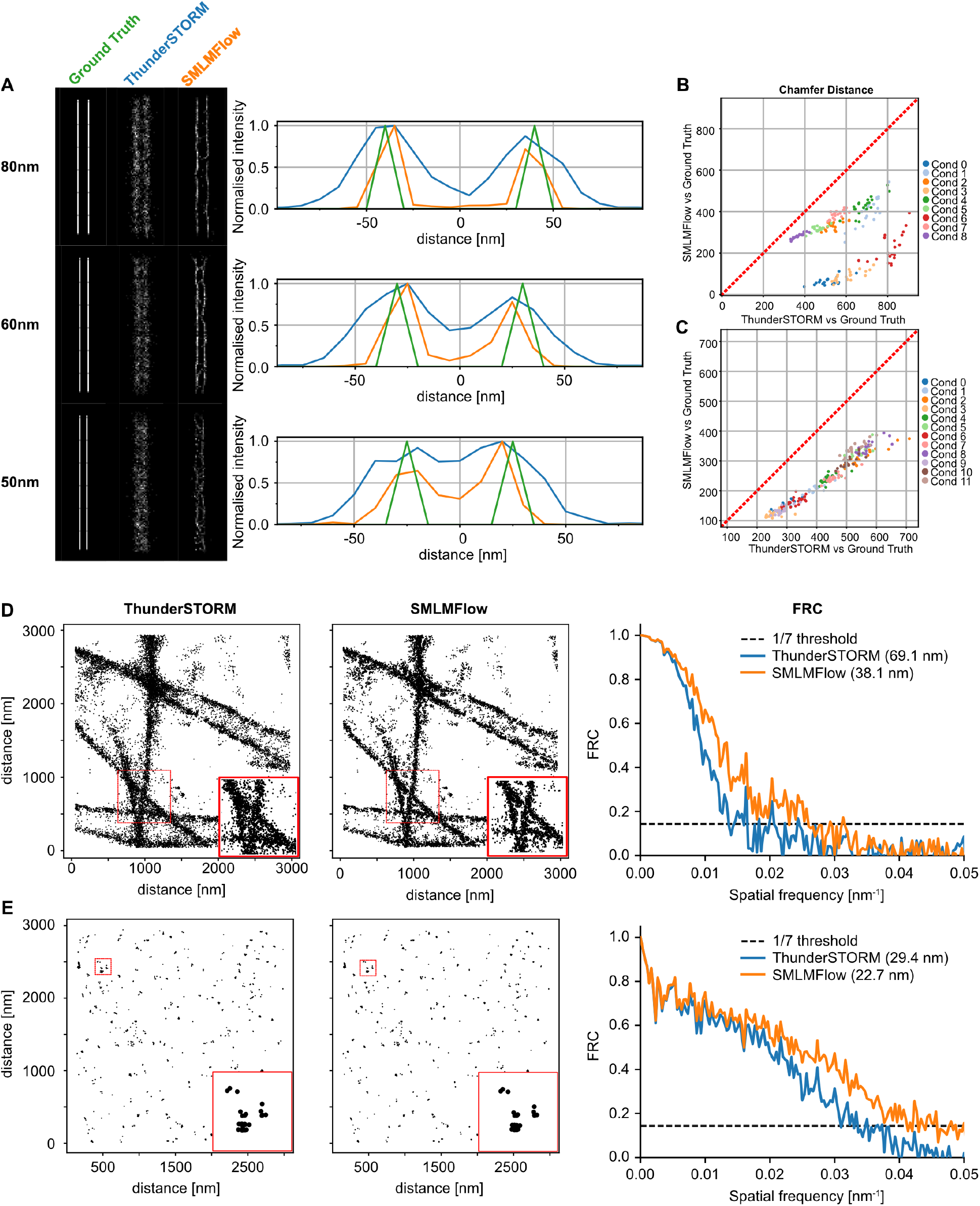
**(A)** Quantitative analysis of SMLMFlow’s structural resolution improvement on synthetic parallel lines. Line plots indicate a separation of the two lines at 50 nm with SMLMFlow, while the initial ThunderSTORM localizations fail after 60 nm. **(B)** Chamfer distance evaluation on the simulated fibrous unknown photophysics dataset. **(C)** Chamfer distance evaluation on the simulated nano-cluster unknown photophysics dataset. **(D)** Qualitative and quantitative evaluation of SMLMFlow on an experimental microtubule dataset. **(E)** Qualitative and quantitative evaluation of SMLMFlow on an experimental protein nano-cluster dataset.

**Figure 3.**
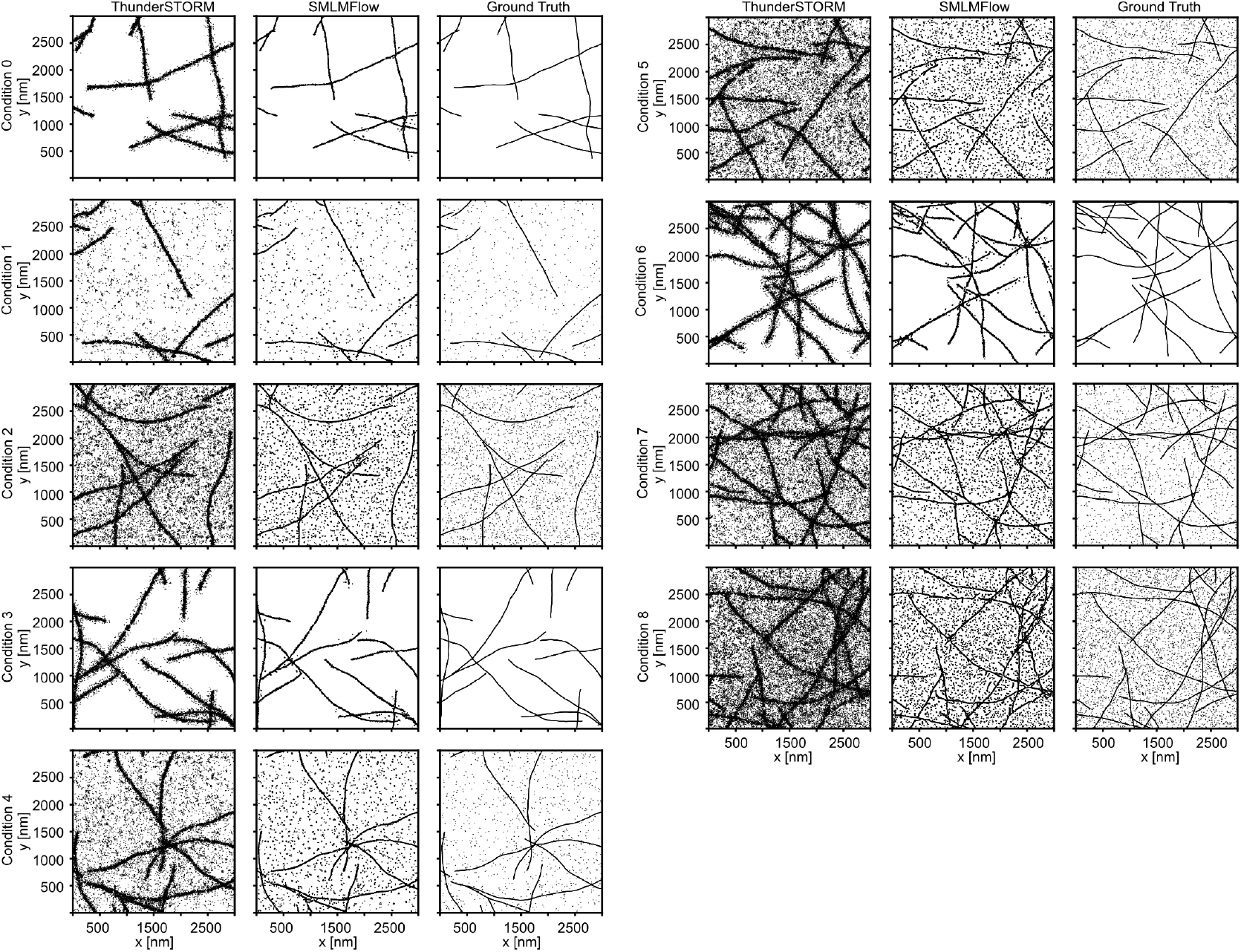
Example result of each simulated fibrous condition.

**Figure 4.**
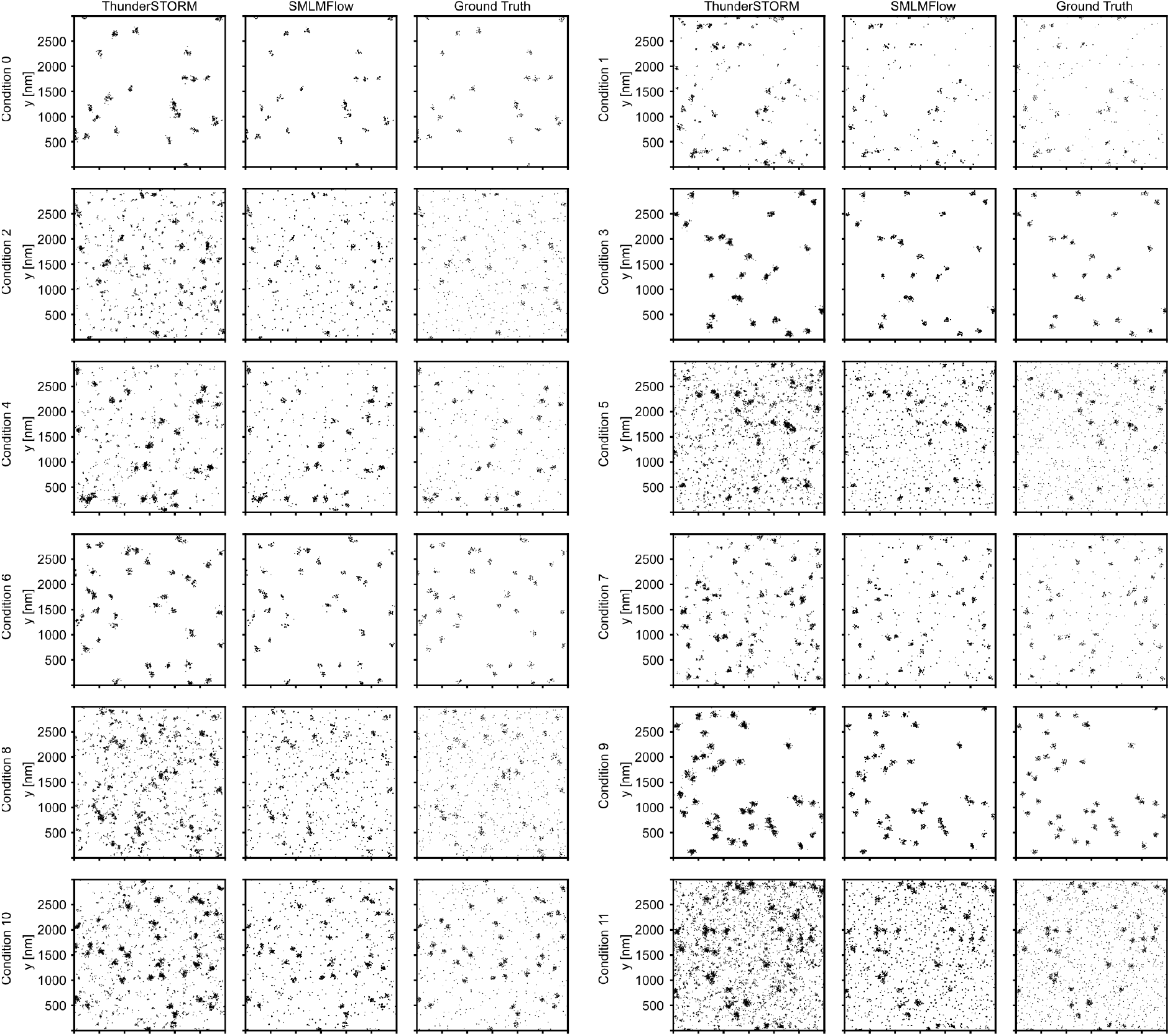
Example result of each simulated nano-cluster condition

**Figure 5.**
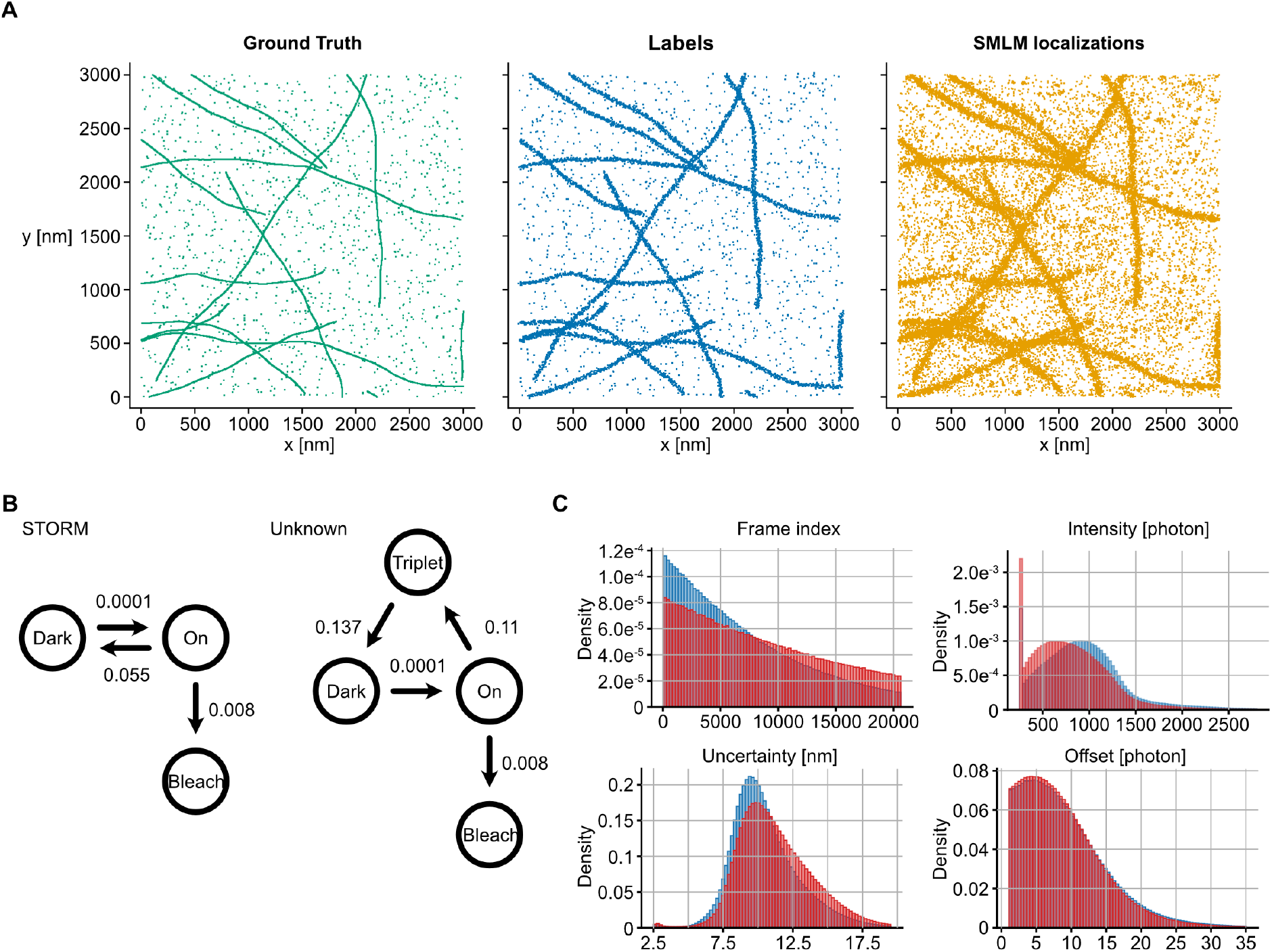
**(A)** Example of a simulated point cloud through the vSMLM process. **(B)** The two photophysics models used to simulate the datasets. **(C)** Histogram comparison of the features of the fibrous dataset between the two photophysics models (blue: STORM; red: Unknown)

**Figure 6.**
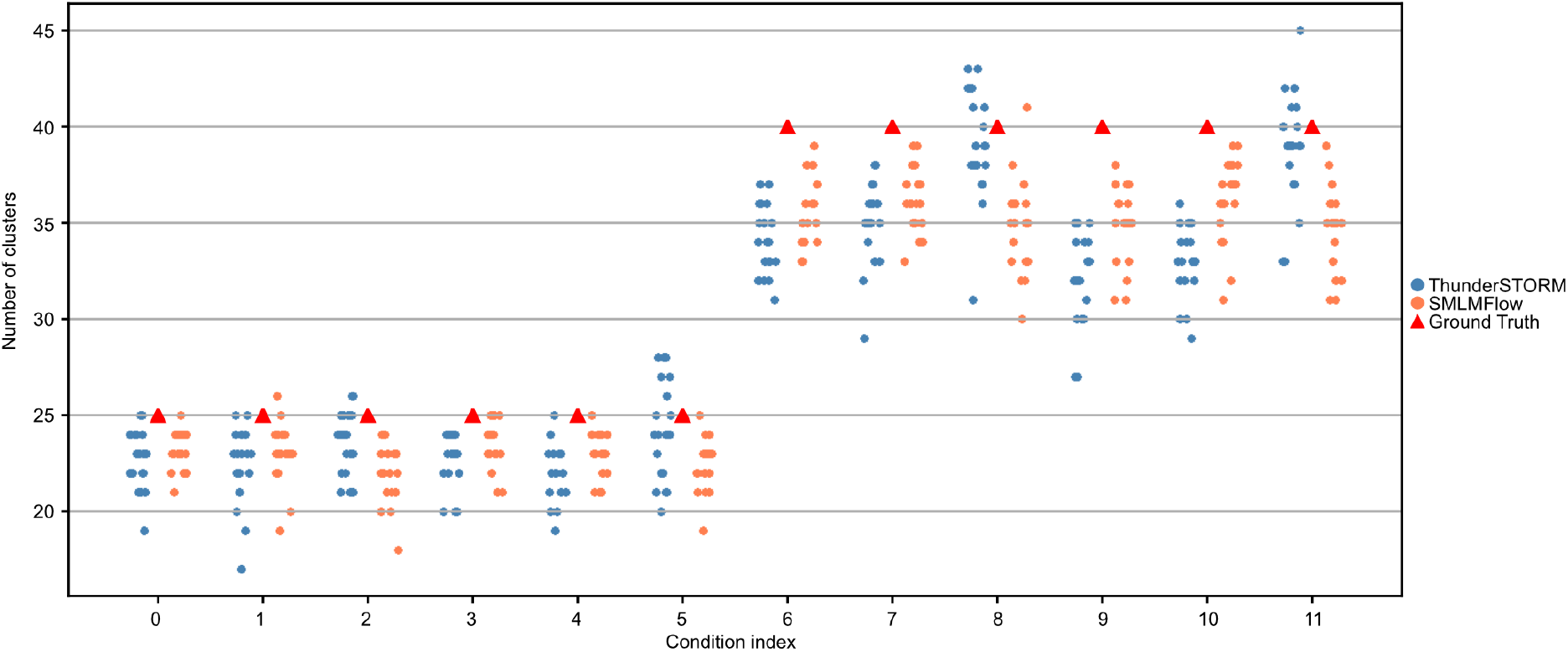
“Number of cluster” evaluation of SMLMFlow vs. ThunderSTORM in the simulated STORM dataset

We next assessed the model’s generalization. Despite being trained under a three-state model, SMLMFlow improved reconstruction on data simulated with an unseen STORM four-state photophysics model (Fig. 2B, C), demonstrating robustness to varying and new imaging conditions. Finally, application to experimental datasets, including microtubules (Fig. 2D) and clustered samples (Fig. 2E), confirmed robust performance on complex, noisy data, with consistent improvements in structural resolution (31 nm for microtubules and 6.7 nm for nano-clusters), as measured quantitatively through the Fourier Ring Correlation (FRC) [18], [19].

## Discussion

In this work, we present SMLMFlow, a novel framework for improving structural resolution in SMLM datasets by directly operating on localization point clouds. Across a broad range of simulated conditions with available ground truth and experimental conditions, SMLMFlow consistently improves structural resolution, remaining robust to variations in noise, sampling density, and photophysics models. Importantly, the ability to generalize to unseen conditions indicates that the model captures the underlying structure of the SMLM data. SMLMFlow is designed to extend existing localization algorithms and can be readily integrated into current SMLM workflows as well as to process already existing SMLM datasets. More generally, it highlights the potential of generative, point cloud-based approaches to recover biologically meaningful structures from noisy data and to bridge the gap between localization precision and true structural resolution.

## ACKNOWLEDGEMENTS

The authors thank Vsevolod Viliuga and Matteo Tadiello for valuable discussions that helped shape the model’s architecture.

## AUTHOR CONTRIBUTIONS

J.G. supervised the project. S.B. and J.G. designed the project. R.P. acquired the experimental data. S.B. implemented, tested and optimised the code. S.B and I.C. ran quantification. J.G., D.S., L.P. and S.B. implemented the simulations. S.B. and J.G. wrote the manuscript. S.B., I.C. and E.L. produced figures. All authors revised the manuscript.

## DATA AVAILABILITY

The data will be made publicly available upon publication.

## CODE AVAILABILITY

All code developed in this manuscript can be accessed through a GitHub repository, which will be made public upon publication.

## Online Methods

### Data (Simulation)

Training a supervised model to improve structural resolution in SMLM data requires paired examples of ground truth coordinates and their corresponding fitted localizations. Such pairs are inherently unavailable in experimental SMLM, where the true fluorophore positions cannot be measured. We therefore relied entirely on synthetic training data, generated using a custom Python-based adaptation of the Virtual-SMLM simulation framework [1]. This framework reproduces the principal sources of noise and degradation in the SMLM acquisition pipeline, including fluorophore photophysics, optical image formation, and camera noise, while retaining access to the underlying ground-truth coordinates. We refer to the original publication for a detailed description of the simulator and focus here on the choices specific to this work.

Two classes of synthetic structures were generated to represent distinct spatial organizations.

#### Filaments

Filamentous structures were simulated using a worm-like chain model with a persistence length of 17 μm and a step size of 3.4 nm between consecutive points, in line with reported biophysical properties of cellular fibers [2]. Multiple fibers were placed within a 3,000×3,000 nm field of view, with the number of filaments and the ratio of uniformly distributed background points varied across nine conditions spanning a range of densities and signal-to-background regimes (Online Methods Table 1 and Online Methods Fig.3.

**Table 1.** Filaments conditions.

| Condition index | Number of fibers | Background ratio [%] |
| --- | --- | --- |
| 0 | 10 | 0 |
| 1 | 10 | 15 |
| 2 | 10 | 33 |
| 3 | 15 | 0 |
| 4 | 15 | 15 |
| 5 | 15 | 33 |
| 6 | 25 | 0 |
| 7 | 25 | 15 |
| 8 | 25 | 33 |

#### Gaussian clusters

Nano-cluster structures were generated by sampling points from Gaussian distributions with a fixed standard deviation of 25 nm. The number of clusters, the number of points per cluster, and the background point ratio were varied across twelve conditions, covering a range of clustering scenarios and noise levels (Online Methods Table 2 and Online Methods Fig.4).

**Table 2.** Gaussian cluster conditions.

| Condition Index | Number of clusters | Number of points per clusters | Background ratio [%] |
| --- | --- | --- | --- |
| 0 | 25 | 15 | 0 |
| 1 | 25 | 15 | 25 |
| 2 | 25 | 15 | 50 |
| 3 | 25 | 30 | 0 |
| 4 | 25 | 30 | 25 |
| 5 | 25 | 30 | 50 |
| 6 | 40 | 15 | 0 |
| 7 | 40 | 15 | 25 |
| 8 | 40 | 15 | 50 |
| 9 | 40 | 30 | 0 |
| 10 | 40 | 30 | 25 |
| 11 | 40 | 30 | 50 |

#### Simulation protocol

For each condition, 100 independent ground-truth structures were generated and then simulated using a three-state STORM photophysical model with dark, on, and bleached states (Online Methods Fig.5B, left). Of these, 80 samples were used for training, and 20 were held out for validation. To evaluate generalization to unseen acquisition conditions, the same 20 validation structures were re-simulated using an alternative photophysical model that extends the standard scheme by adding a triplet state between the on and dark states (Online Methods Fig.5B,right). This addition produces more complex blinking dynamics and represents a distributional shift relative to the STORM model (Online Methods Fig.5C). These samples were used exclusively to test the model’s robustness to previously unseen photophysical behavior.

#### Labeling model

All simulations used an immunostaining-based labeling strategy with fixed parameters across conditions. A primary antibody was applied with full labeling efficiency, followed by a secondary antibody with a one-to-one binding constraint. Spatial offsets corresponding to antibody sizes were incorporated to reflect realistic localization displacements introduced during labeling (Table 3).

**Table 3.**
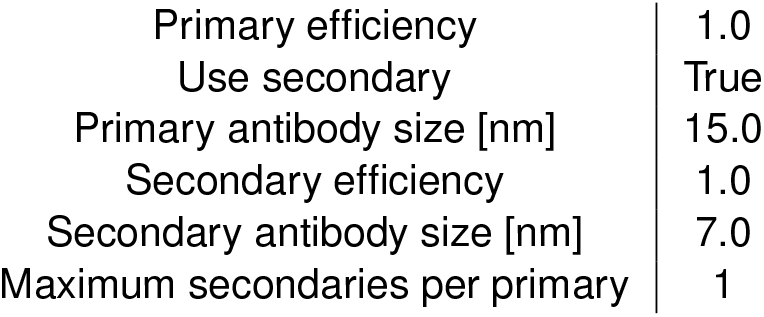
Labeling model parameters.

|  |  |
| --- | --- |
| Primary efficiency | 1.0 |
| Use secondary | True |
| Primary antibody size [nm] | 15.0 |
| Secondary efficiency | 1.0 |
| Secondary antibody size [nm] | 7.0 |
| Maximum secondaries per primary | 1 |

#### Camera model

Simulated image acquisition used a fixed camera configuration representative of an EMCCD system, with a 100 Hz frame rate, 100 nm pixel size, quantum efficiency of 0.9, and realistic noise sources including readout noise, spurious charge, and electron-multiplication gain. The field of view was pixelated into a 50 × 50-pixel grid (Table 4).

**Table 4.** Camera model parameters.

|  |  |
| --- | --- |
| Framerate [Hz] | 100.0 |
| X pixel size [nm] | 100.0 |
| Y pixel size [nm] | 100.0 |
| X size [pixels] | 50 |
| Y size [pixels] | 50 |
| Quantum efficiency | 0.9 |
| Readout noise [$e^-$] | 74.4 |
| Spurious charge [$e^-$/pixel/frame] | 0.0002 |
| EM gain | 300.0 |
| Baseline [ADU] | 100.0 |
| Electrons per ADU [$e^-$/ADU] | 45.0 |
| Bit depth [bits] | 16 |
| Autofluorescence [photons] | 5.0 |

#### Fluorophore model

Fluorophore emission was simulated using a physically motivated model that incorporated photon emission rates, collection efficiency, and point-spread function characteristics. We chose emission properties to reflect an Alexa Fluor 647 dye with realistic photon budgets and noise levels (Table 5) [3].

**Table 5.** Fluorophore model parameters.

|  |  |
| --- | --- |
| Emission rate [photons/s] | 78870 |
| Collection efficiency | 0.42 |
| Z factor | 2.5 |
| PSF FWHM [nm] | 352.5 |
| Background noise [photons] | 100 |
| Excitation wavelength [nm] | 647 |
| Extinction coefficient [$M^{-1}cm^{-1}$] | 239000 |
| Quantum yield | 0.33 |
| Excitation intensity [$W/cm^2$] | 100000 |

**Table 6.** Configuration of the ERWIN Transformer.

|  |  |
| --- | --- |
| Feature dimension | 64 |
| Encoder depths | [2,2,2,6,2] |
| Decoder depths | [2,2,2,2] |
| Encoder attention heads per level | [2,4,8,16,32] |
| Decoder attention heads per level | [4,4,8,16] |
| Hierarchical strides | [2,2,2,2] |
| Ball sizes | [256,256,256,128,64] |
| Initial message passing steps | 3 |

#### Localization extraction

Simulated image sequences were processed using a standard SMLM localization pipeline. Specifically, we applied the ThunderSTORM multi-emitter fitting algorithm [4] to detect and localize fluorophores from the simulated frames (Fig.5A).

#### Output data

For each simulation, the pipeline produced paired data consisting of ground truth molecular coordinates and corresponding noisy localization coordinates obtained from ThunderSTORM in an Multi Emitter Fitting (MEF) scheme with the emitter setting of 3. These paired datasets were used to train SMLMFlow.

### Data (Real)

#### Protein clusters

Clustered protein distributions were obtained from the publicly available nano-org resource [5], a curated database of SMLM datasets. From this repository, we selected a representative STORM dataset of Lck on activated T cells (file: `20260305_Lck_aCD3CD28_region0.csv`) and extracted a single region of interest for evaluation, measuring 3,000 nm × 3,000 nm.

#### Microtubules

The microtubules dataset was also obtained from the nano-org resource with the file name `Sandeep_microtubules_cos7_CF660_cell-3_filtered_merged.csv`, and cut to a 3,000nm × 3,000nm region of interest.

### Evaluation

#### Chamfer Distance

To quantitatively assess the structural accuracy of reconstructed point clouds, we use a one-directional Chamfer distance from source to ground truth as our primary evaluation metric [6]. The Chamfer distance is well-suited to SMLM evaluation because both the predicted and ground-truth data are unordered point clouds with potentially different cardinalities, and classical metrics that rely on pointwise correspondence or ordering are therefore not applicable.

For two point clouds *X* = {*x*_*i*_} and *Y* = {*y*_*i*_}, the bidirectional Chamfer distance *CD*_*bi*_ is defined as

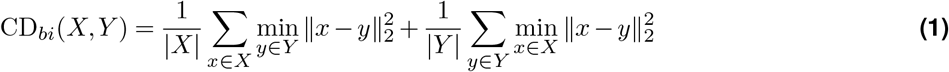

In our setting, however, the source-to-target direction is the quantity of biological interest: it asks whether each predicted localization lies close to a true emitter, which is the correct measure of whether the reconstructed structure faithfully represents the underlying biology. The reverse direction, ground truth-to-source, is dominated by background and labeling stochasticity rather than reconstruction quality. In particular, ground truth points that are unlabeled (e.g., due to imperfect labeling efficiency) or undersampled by ThunderSTORM contribute large nearest neighbor distances that do not reflect any failure of SMLMFlow itself, and that are present in both the input ThunderSTORM localizations and the SMLMFlow output to a similar degree. Including this term, therefore, inflates the metric with sampling artifacts common to all methods and obscures genuine differences in reconstruction.

We therefore report the one-directional source-to-target Chamfer distance *CD*_*st*_,

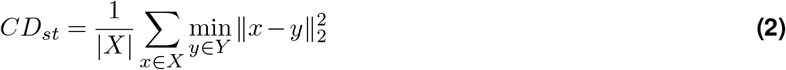

Where X is the reconstructed point cloud (either ThunderSTORM output or SMLMFlow output), and Y is the ground truth. This measures, for each predicted point, the distance to its nearest ground-truth emitter, providing a direct quantitative measure of how closely the reconstructed localizations align with the underlying structure.

#### Fourier Ring Correlation

As a complementary, image-based measure of effective resolution, we report the Fourier Ring Correlation (FRC), a standard metric for quantifying resolution in single-molecule localization microscopy [7]. Unlike the Chamfer distance, which operates directly on point coordinates and requires access to ground-truth emitter positions, FRC is computed from rendered images and does not depend on ground truth, making it applicable to both simulated and experimental data.

FRC quantifies resolution by measuring the spatial-frequency content shared between two independent reconstructions of the same structure. The localizations of a given region of interest are randomly partitioned into two statistically independent halves, which are rendered into two images *I*_1_ and *I*_2_ at a fixed pixel size. The normalized cross-correlation between their Fourier transforms is then computed over rings of constant spatial frequency *q*,

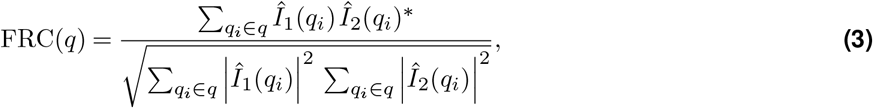

where *Î*_1_ and *Î*_2_ denote the discrete Fourier transforms of the two half-images, ^∗^ denotes complex conjugation, and the sum runs over all frequencies *q*_*i*_ falling within the ring of radius *q*. The FRC curve decays from unity at low frequencies, where both images are dominated by correlated signal, toward zero at high frequencies, where they are dominated by uncorrelated noise. The effective image resolution is defined as the inverse of the spatial frequency *q*_1*/*7_ at which the FRC curve first crosses the fixed threshold of 1*/*7, such that the resolution is given by 1*/q*_1*/*7_.In practice, FRC values were computed using the NanoPyx implementation [8]. For each region of interest, localizations were split into two independent subsets (odd and even frame indices), rendered onto a common pixel grid, and processed with the NanoPyx FRC routine to obtain the resolution estimate (Fig. 2D, E).

### Flow Matching

Flow matching is a simulation-free method for training continuous normalizing flows (CNFs) [9]. Given a source distribution *p*_0_ and a target data distribution *p*_1_, the goal is to learn a time-dependent velocity field *v*_*θ*_(*x, t*), *t*∈ [0, 1], that transports samples from *p*_0_ to *p*_1_ by defining an ordinary differential equation (ODE):

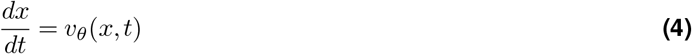

whose solution yields the flow map

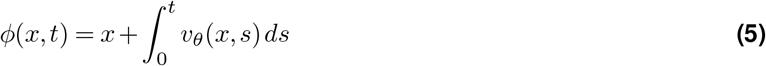

A probability path *p*_*t*_ interpolating between *p*_0_ and *p*_1_ is constructed via per-point linear interpolation between paired samples *x*_0_ ~ *p*_0_ and *x*_1_ ~ *p*_1_:

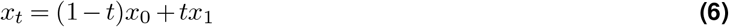

The target velocity at each interpolated point is then *u*_*t*_(*x*_*t*_) = *x*_1_ − *x*_0_. The model is trained by minimizing the mean squared error between the predicted and target velocities:

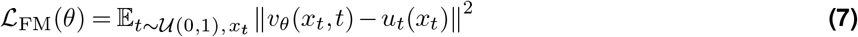

In contrast to diffusion models [10], which require iterative stochastic denoising along a fixed Gaussian noise schedule, flow matching enables efficient and deterministic generation by integrating a single learned ODE [9; 11].

#### Conditional Flow Matching

In conditional flow matching (CFM) [9], the velocity field is additionally conditioned on a context variable *c*, yielding *v*_*θ*_(*x, t, c*). Rather than learning a single global transport between *p*_0_ and *p*_1_, CFM constructs the probability path and target velocity conditioned on individual data samples, making the training objective tractable. Specifically, conditioning on a target sample *x*_1_ defines a per-sample probability path *p*_*t*_(*x* |*x*_1_) with a closed-form target velocity *u*_*t*_(*x* |*x*_1_) = *x*_1_−*x*_0_, whose expectation over *p*_1_ recovers the marginal flow [9; 11].

In our setting, the conditioning variable *c* encodes structural context derived from the input localization point cloud: local molecular neighborhoods and extended structural patterns such as fiber orientation and cluster geometry, as extracted by the Erwin transformer. This is closely related to the meta flow matching framework [12], which conditions the velocity field on task-level context to generalize across heterogeneous distributions. Here, structural context conditioning allows the model to adapt its transport to the local geometry of each ROI, rather than learning a single average transformation across all localization patterns.

#### Application to SMLM Point Clouds

A central challenge in applying flow matching to SMLM data is that source localizations and ground-truth emitter coordinates are inherently unpaired. A single ROI typically contains a different number of reconstructed localizations than ground-truth points, and no one-to-one correspondence between them exists. Standard conditional flow matching either assumes paired samples or learns a stochastic coupling between the two distributions [11], neither of which is directly applicable here.

To define a tractable supervision target on point clouds of differing cardinality, we construct a deterministic source-to-target coupling via nearest-neighbor assignment. For each source point *x* ∈ *X*_0_, we assign a target:

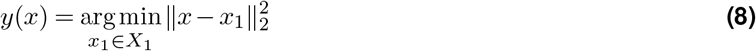

This assignment is computed once per sample at the start of training and held fixed thereafter, providing each source point with a stable per-point regression target. The assignment is one-directional and not bijective: multiple source points may map to the same target, while some ground-truth points may remain unassigned. We found this simple coupling sufficient for our setting.

With this assignment in place, let *X*_0_ ~*p*_0_ denote the SMLM localizations and *X*_1_ ={*y*(*x*) : *x*∈ *X*_0_} the corresponding nearest-neighbor assigned ground-truth points. Training reduces to the standard conditional flow matching objective, with the velocity field conditioned on the structural context *c* extracted from *X*_0_:

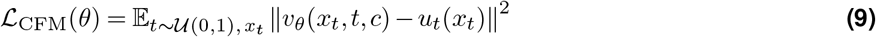

computed per-point and averaged across all points in each ROI and across the batch.

At inference time, structural reconstruction is performed by solving the learned ODE from the observed noisy localization point cloud using an Euler ODE solver over *N* steps, where *N* is a user-defined parameter:

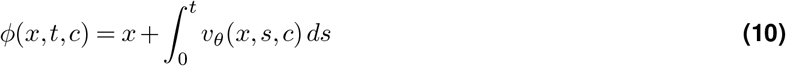

During development, we observed that reconstruction quality does not necessarily peak at the trajectory endpoint *t* = 1. Instead, integrating the full flow can over-sharpen the reconstruction and introduce structural artifacts, manifesting as a degradation in the Chamfer distance. The optimal stopping point is data-dependent: on simulated data, reconstruction quality was highest at approximately *t* = 0.8, whereas on experimental data the optimum occurred substantially earlier, around *t* = 0.2. We attribute this difference to the distributional gap between simulated and experimental localizations, and therefore treat the integration endpoint as a user-tunable parameter, reporting results at the empirically optimal stopping time for each dataset.

### Model

SMLMFlow is a neural network that predicts the time-dependent conditional velocity field *v*_*θ*_(*x*_*t*_, *t, c*) underlying the flow matching objective described above. It operates directly on unordered point clouds and outputs a per-point 2D displacement vector conditioned on both spatial context and acquisition-specific features. The architecture comprises three main components: an SMLM feature embedding, a hierarchical transformer based on the Erwin transformer [13], and a flow decoding head.

#### SMLM feature embedding

Each localization is associated with experimentally derived attributes: frame index, photon count, localization uncertainty, and background intensity. Frame indices are encoded using sinusoidal positional embeddings into a 64-dimensional vector. Photon count and background intensity are passed through a log(1+x) transform for numerical stability, and then combined to form a per-point signal-to-noise ratio. Both this signal-to-noise ratio and the localization uncertainty are mapped into 64-dimensional vectors via small two-layer MLPs with GELU activations. The three resulting embeddings are concatenated and projected back to 64 dimensions through a final two-layer MLP, yielding a single per-point feature vector that serves as the input to the backbone. Spatial coordinates themselves are not part of the feature embedding; they enter the model exclusively as point positions used by the transformer’s ball-tree partitioning and message passing.

#### Graph construction and local geometric encoding

To incorporate local geometric context, a graph is constructed dynamically for each input point cloud based on Euclidean proximity. We construct a k-nearest-neighbor graph with k = 10 and additionally remove edges longer than 300 nm. The kNN construction bounds the node degree, keeping the edge count linear in the number of localizations (10N) and the memory and message-passing cost predictable despite the strong spatial density variations characteristic of SMLM data. The 300 nm cutoff enforces physical locality: it removes the long, spurious edges that kNN can introduce in sparse regions, restricting message passing to localizations within a length scale over which they plausibly belong to the same underlying structure. Together, the two criteria yield a graph that is both computationally bounded in dense regions and physically meaningful in sparse ones. This graph is consumed by the first block of the Erwin backbone, which applies three steps of relative-position-conditioned message passing before the hierarchical attention layers. This initial message passing provides a short-range geometric prior that complements the longer-range context captured by the subsequent ball-tree attention.

#### Hierarchical transformer backbone

Spatial feature extraction is performed using the Erwin transformer [13]. We adopt the original architecture without modification and refer the reader to that work for full details. The point cloud is recursively partitioned into local neighborhoods (“balls”) using a ball-tree structure, and within each ball, multi-head self-attention is applied to capture local spatial dependencies. A hierarchical encoder–decoder structure aggregates information across scales using ball-based pooling and unpooling operations. To improve robustness to point ordering, the model employs deterministic ball-tree reordering with alternating rotations across transformer layers, as implemented in the original architecture. In our implementation, the backbone is configured as follows:

#### Time conditioning

To model the continuous transformation between noisy and ground truth point clouds, the network is conditioned on a time variable *t* ∈ [0, 1]. Time is embedded using a 32-dimensional sinusoidal encoding, concatenated with the per-point spatial features produced by the transformer backbone, and then passed to the flow decoder.

#### Flow decoder

The flow decoder is a multi-layer perceptron that maps the combined spatial and temporal features to a 2D displacement vector for each point. It consists of four fully connected layers with hidden dimension 256, GELU activations, and a final linear projection to a 2D output that constitutes the predicted velocity field *v*_*θ*_(*x*_*t*_, *t, c*).

#### Training details

The model is trained using the flow matching objective described above. For each ROI, a time *t* ~*U* (0, 1) is sampled, intermediate states are constructed via linear interpolation between the source and its assigned ground truth points, and the model predicts the corresponding velocity field. The loss is the mean squared error between predicted and target velocities, averaged across all points in each ROI and across the batch. Optimization is performed using Adam with a learning rate of 5·10^*−*4^. Models are trained for 400 epochs on 4 GPUs with an effective batch size of 8.

#### Inference

At inference time, the learned velocity field is integrated from the observed noisy localization point cloud using a forward-Euler scheme with 20 steps. As described in the Flow Matching section, integration is terminated early depending on the specific dataset.

### Further Experiments

#### Cluster descriptor recovery on simulated data

To assess whether SMLMFlow’s improvements in structural reconstruction translate into more accurate downstream biological quantification, we performed a clustering analysis on the simulated Gaussian cluster dataset. We compared the recovered cluster descriptors against the known ground truth. Both ThunderSTORM and SMLMFlow outputs were processed through the same analysis pipeline, applied identically to each. Cluster identification was performed using ToMATo, a topological clustering algorithm that combines a density estimate with persistent homology to identify modes in the point distribution and merge those whose persistence falls below a chosen threshold [14]. We used a logarithmic distance-to-measure density estimate with a per-sample neighborhood graph radius automatically set to the argmax of Ripley’s H-function, capped at 200 nm. The persistence threshold and minimum cluster size were derived from the known generative parameters of each condition — the expected number of clusters n, points per cluster p, and background ratio b — and scaled by the ratio of observed to expected total points to account for variations in localization yield. Clusters smaller than the resulting minimum size were assigned to a noise label and excluded from descriptor computation. For each retained cluster, we computed an effective spatial extent *σ*_*cluster*_ = *mean*(*λ*_*i*_) where *λ*_*i*_ are the eigenvalues of the per-cluster spatial covariance matrix. This isotropic-equivalent standard deviation is directly comparable to the ground truth value of 25 nm used during simulation.

Across all twelve cluster conditions, descriptors derived from SMLMFlow-processed data more closely matched the ground truth than those derived from ThunderSTORM. The recovered per-cluster size *σ*_*cluster*_ converged toward the simulated 25 nm value (Fig. 1D), and the recovered number of clusters stayed consistent between ThunderSTORM and SMLMFlow across conditions (Online Methods Fig. 6). Together, these results show that the structural improvements demonstrated by Chamfer distance translate into more accurate biological quantification on standard downstream analyses.

#### Resolution evaluation on parallel-lines

To quantify the effective structural resolution of SMLMFlow, we generated a dataset of pairs of parallel lines separated by varying distances. We processed each through the full simulation, reconstruction, and SMLMFlow pipeline. Each sample consisted of two infinitesimally thin parallel lines, each 1,000 nm long, separated by a gap ranging from 30 nm to 150 nm. The simulation, labeling, and acquisition pipeline were identical to those used for the filamentous training data, and ThunderSTORM was used to obtain the corresponding noisy reconstructions. As complementary baselines, eSRRF [15] from NanoPyx [8] was applied to the same simulated image stacks to provide a non-localization-based super-resolution reference. For all conditions, a ring radius of 2.0, sensitivity of 3, and eSRRF order of 3 were used, as they were the best-performing parameters found. SMLMFlow predictions were obtained from the filamentous-trained model with no fine-tuning on the parallel-line geometry, providing a strict out-of-distribution evaluation. To enable direct comparison between methods, all point clouds were rendered into binned 2D images at a fixed pixel size of 10 nm. The y-axis range was set globally across all samples to ensure a consistent reference frame, while the x-axis range was determined per sample based on the data’s spatial extent, with a small constant padding. Each binned image was normalized by its peak intensity. eSRRF reconstructions, produced at 20 nm/pixel, were resampled and rescaled to match the 10 nm/pixel grid used by the localization-based methods. To remove residual translational offsets between methods and across samples, all images were aligned to a common structural center derived from the ground truth. The final intensity line profiles shown in Fig. 2A and Online Methods Fig. 7 were extracted from the aligned images by averaging across the full fibers’ length in the y direction. This procedure was repeated for line separations of 30, 40, 50, 60, 80, 100, 120, and 150 nm. The minimum separation at which a method produces two visually resolvable peaks defines its effective structural resolution. We observe a 10nm improvement between ThunderSTORM and SMLMFlow for 50nm and 60nm-spaced parallel lines.

**Figure 7.**
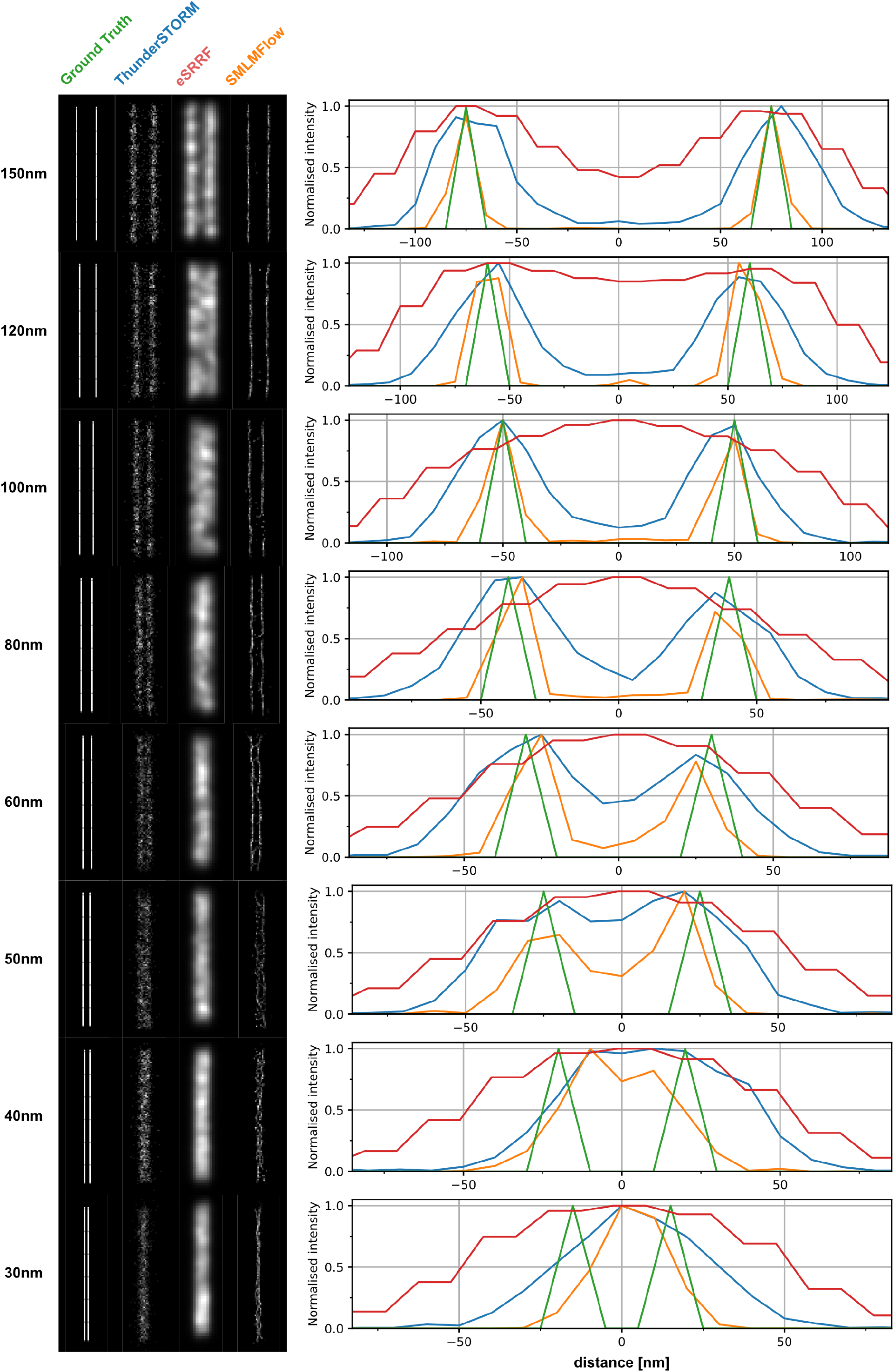
Parallel lines evaluation between ThunderSTORM, eSRRF, and SMLMFlow ranging from 150nm to 30nm.

